# Structural basis of coiled-coil domain-mediated inhibition of *A. baumannii* AbTir NAD^+^ nucleosidase activity

**DOI:** 10.64898/2026.10.06.757174

**Authors:** Weixi Gu, Mohammad K. Manik, Chacko Jobichen, Matthieu Haudiquet, Jovita D’Silva, Chen-Yang Lai, Sulin Li, Xiaoqi Qian, Ashar J. Malik, David B. Ascher, Randy J. Read, Jeremy J. Barr, Wai-Hong Tham, Aude Bernheim, Gavin J. Knott, Bostjan Kobe

## Abstract

TIR (Toll/interleukin-1 receptor) domains are central components of innate immunity and cell-death pathways across diverse forms of life. In bacteria, TIR domain-containing proteins participate in diverse antiphage defence systems, several of which rely on NADase activity carried out by TIR domains. This nucleosidase activity is often regulated by additional domains that modulate activation and specificity in response to infection. AbTir, a TIR domain-containing protein from *Acinetobacter baumannii*, contains an N-terminal CC domain that negatively regulates its NADase activity, but the structural basis of this regulation remains unknown. Here, we used X-ray crystallography to investigate the molecular basis of CC domain-mediated regulation in AbTir. We determined the crystal structure of full-length AbTir in complex with a nanobody, revealing a symmetric AbTir TIR dimer in which only one of the two symmetry-related CC helices is accommodated at the dimer surface, breaking the overall symmetry of the full-length protein. Structural comparisons with other bacterial TIR domains suggest that such TIR dimers likely represent an inactive state, in which the NAD^+^-binding site has not yet formed. Phylogenetic analysis places AbTir within a family of bacterial TIR proteins associated with defence-related genomic loci, although phage-spot assays performed in the surrogate host *Escherichia coli* did not detect measurable antiphage activity. Together, these findings reveal an unusual structural mechanism of CC domain-mediated inhibition of a bacterial TIR NADase and expand our understanding of the mechanistic diversity of bacterial TIR proteins.

## Introduction

Toll/interleukin-1 receptor (TIR) domains are cytoplasmic domains found in proteins from bacteria, archaea, plants and animals^1^. They share a conserved flavodoxin-like fold with a five-stranded β-sheet core (βA-βE) surrounded by five α-helices (αA-αE)^1^. TIR domains typically play roles in innate immunity and cell-death pathways^1–3^; and generally function through self-association or association with other TIR domains, forming filamentous assemblies^4,5^.

In bacteria, TIR domain-containing proteins (Tcps) were first identified as virulence factors that enable pathogenic bacteria to evade and suppress the host innate immunity^6^. They exert this function through interactions with host TIR domain-containing proteins such as Toll-like receptors (TLRs) and their adaptor proteins, disrupting signalling pathways in a process termed “molecular mimicry”^7^. More recently, Tcps have been found to catalyse the cleavage of NAD^+^ into nicotinamide and ADPR (ADP-ribose) or different cyclic ADPR variants (cADPR, 2’cADPR, 3’cADPR and N7-cADPR)^8–10^, and base-exchange products such as imidazole-ADPR and His-ADPR^11,12^. Tcps are now recognised as widespread components of bacterial antiphage systems^13,14^, including the Thoeris defence system^15^ and cyclic GMP-AMP synthase (cGAS)-stimulator of interferon genes (STING) pathway^16^, in which their NAD^+^-cleavage activity is essential.

A general feature of TIR-domain function appears to be a head-to-tail association of TIR domains into filamentous assemblies^4,5^, through a structural motif called the “BB-loop” (between the βB strand and αB helix). This head-to-tail self-association is essential for NADase activity, because the active site forms in the corresponding head-to-tail interface. The formation of filamentous signalosomes leads to specific properties of the corresponding signalling mechanism termed SCAF (signalling by cooperative assembly formation)^3,17,18^.

The activity of bacterial Tcps is typically regulated by their diverse accessory domains, which provide distinct mechanisms of regulation^13,19^. For example, many bacterial Tcps, including TcpB from *Brucella melitensis*, TcpS from *Salmonella* Enteritidis and TirS from *Staphylococcus aureus*, possess a coiled-coil (CC) domain (Figure S1A). In several cases, these CC regions have been shown to regulate TIR-domain function and contribute to immune modulation^20–23^. CC domains are ubiquitous domains found in all phyla, and are best known for mediating protein oligomerisation^24,25^. They often serve as molecular spacers that define the distance between protein modules, and can also function as membrane-anchoring elements involved in vesicle tethering and pore formation^24^. Previous studies on TcpB have suggested that its CC domain is conformationally dynamic in solution^22^ and mediates phosphoinositide interaction through a conserved ^46^KKRxxxxKK^54^ motif (Figure S1A), which contributes to the suppression of TLR-mediated innate immune responses^26^.

*Acinetobacter baumannii* is a Gram-negative bacterium that causes opportunistic infections of blood, urinary tract, lungs and skin^27^. Previous studies on the TIR domain-containing protein in *A. baumannii* (AbTir) showed that its C-terminal TIR domain (AbTir^TIR^) hydrolyses NAD^+^, while the N-terminal CC domain (AbTir^CC^) is required for protein dimerization, and negative regulation of this nucleosidase activity^8^. Activation of AbTir involves head-to-tail association of TIR domains, accompanied by large conformational rearrangements that generate the catalytic site at the interface between two monomers^8,28^. In addition to cleaving NAD^+^ into ADPR and nicotinamide, the protein also cyclizes ADPR into 2’cADPR^8^. AbTir has served as a model system for identifying products of NAD^+^ cleavage by TIR domains, as well as elucidating the biological function of 2’cADPR^8,9,29,30^. What remains unclear is the molecular basis of CC domain-mediated inhibition and the physiological role of AbTir.

To investigate the mechanism of CC domain-mediated regulation, we sought to determine the crystal structure of full-length AbTir. As crystallization attempts of either full-length AbTir or isolated AbTir^CC^ were unsuccessful, we generated AbTir-specific nanobodies to facilitate crystallization. Here, we report the crystal structure of full-length AbTir in complex with nanobody A1 (NbA1) at 2.65 Å resolution. Phylogenetic studies place AbTir within a defence-associated family of bacterial TIR proteins, although phage-spot assays performed in the surrogate host *Escherichia coli* did not detect such antiphage activity. Together, this work provides structural insight into how the CC domain stabilizes an inactive conformation and suppresses AbTir nucleosidase activity through an unusual mechanism, while expanding our understanding of the mechanistic diversity of bacterial TIR proteins.

## Results

### Nanobodies modulate AbTir dimerization and inhibit nucleosidase activity

The molecular basis of AbTir NADase activity has previously been characterised^8^. Upon assembly into a filamentous structure, AbTir undergoes large conformational changes that promote formation of the catalytic site at the head-to-tail interface between two protomers. However, the mechanism regulating this transition and the role of the CC domain remain uncharacterized. To address these questions, we sought to determine the structure of full-length AbTir by X-ray crystallography. To overcome the challenges associated with crystallisation of the highly dynamic full-length AbTir, we employed AbTir-specific nanobodies. These single-domain antibodies, known for their high affinity and stability, can act as crystallization chaperones, facilitating crystal formation by stabilizing specific protein conformations and mediating crystal contacts^31^. We generated four distinct AbTir-specific nanobodies, which were classified into two groups (A and B) based on sequence similarities of their complementarity determining regions (CDRs): nanobodies NbA1 and NbA5 in group A; and NbB1 and NbB5 in group B (Figure S1B).

AbTir exhibits concentration-dependent dimerization in solution^8^. To investigate the effects of nanobodies on AbTir, we employed size-exclusion chromatography coupled with multi-angle light scattering (SEC-MALS) to measure the molecular masses of AbTir:nanobody complexes in solution (AbTir: 31 kDa; nanobody: 14 kDa). These experiments yielded molecular masses of 92.7, 102.8, 61.3 and 57.0 kDa for AbTir complexes with NbA1, NbA5, NbB1 and NbB5, respectively (Figure 1A). The results suggest that two class A nanobodies interact with an AbTir dimer, while one class B nanobody binds to an AbTir monomer. We conclude that class A and B nanobodies show different binding modes to the AbTir protein: NbA1 and NbA5 preserve AbTir dimerization, while NbB1 and NbB5 disrupt it.

**Figure 1:**
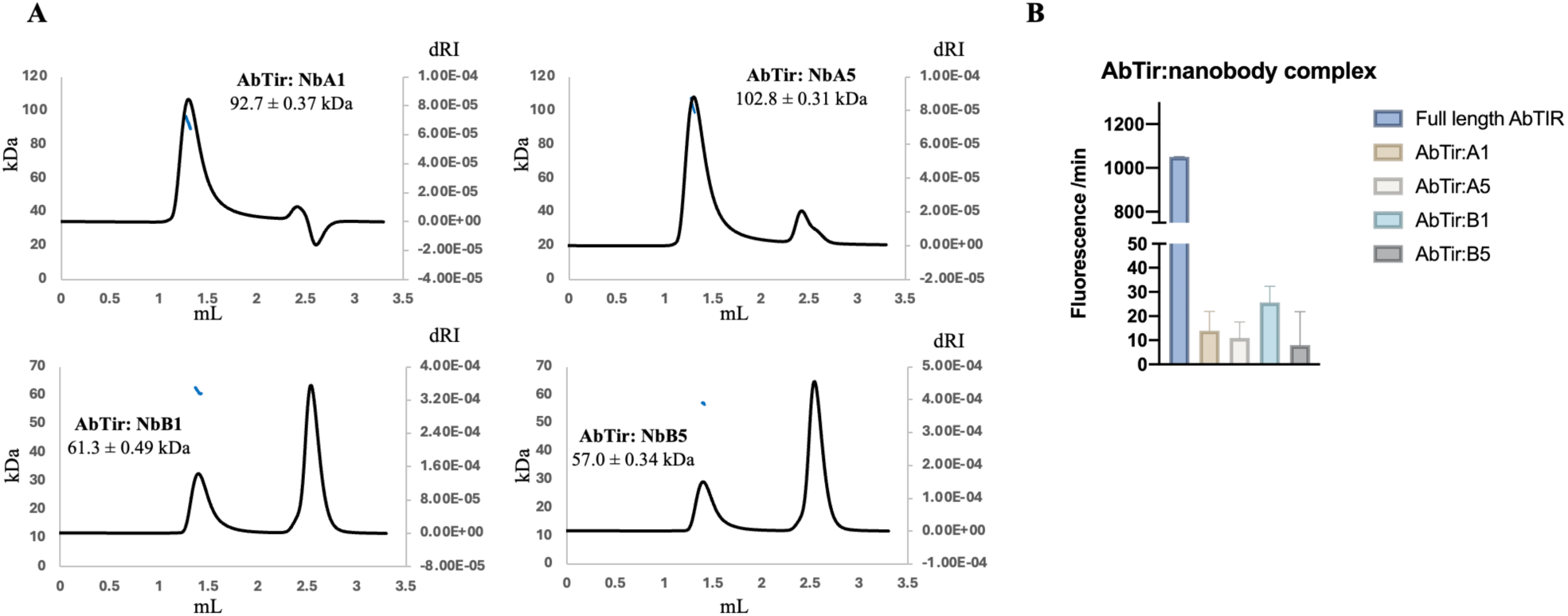
Nanobodies modulate AbTir dimerization and nucleosidase activity. (**A)** SEC-MALS analysis of the AbTir:nanobody complexes. The X-axis shows the elution volume during SEC. The left Y-axis indicates the calculated molecular mass; the solid lines denote the average molecular mass distributions across the peak. The right Y-axis indicates the dRI (differential refractive index) value. (**B)** Fluorescence assays of NADase activity of the AbTir:nanobody complexes.

To investigate the effects of nanobody interaction on AbTir enzymatic activity, we performed fluorescence-based NADase assays. All four AbTir: nanobody complexes showed substantially reduced activity compared with AbTir alone, suggesting that nanobody binding inhibits AbTir NAD^+^ nucleosidase activity (Figure 1B). Given their distinct effects on oligomerization, the two nanobody classes likely inhibit AbTir through different mechanisms. Class A nanobodies may bind to regions critical for catalytic activity, either by blocking the access of the substrate NAD^+^ to the protein, or by stabilizing an inactive, dimeric conformation of the protein. By contrast, class B nanobodies likely inhibit activity by disrupting protein oligomerization required for NAD^+^ cleavage.

### Crystal structure of the NbA1:AbTir complex

To elucidate the mechanistic basis of nanobody-mediated inhibition, we attempted to crystallize the nanobody complexes. Of all the complexes, only the AbTir:NbA1 complex yielded crystals, which diffracted at 2.65 Å resolution. The crystals have the symmetry of the space-group P4_1_ 2 2, with one AbTir and one NbA1 molecule in the asymmetric unit (Figure S2A, Table 1). As the AbTir complex is dimeric in solution, we examined the crystal packing to identify the most likely dimeric assembly (Figure S2A). In the crystal (Figure 2A, S2A), AbTir TIR domains form a symmetric dimer. The long CC helix from each AbTir protomer occupies the same binding surface on the AbTir dimer. These symmetry-related CC helices sterically overlap, indicating that their occupancy is mutually exclusive. The helices from the two protomers of the dimer would occupy the same space, but in opposite directions. The electron density therefore presumably reflects the superposition of the two helices running in opposite directions, consistent with the poor interpretability of the electron density in this region. The side-chain electron densities for the helix were therefore poor (Figure S2B), making residue assignments difficult.

**Figure 2:**
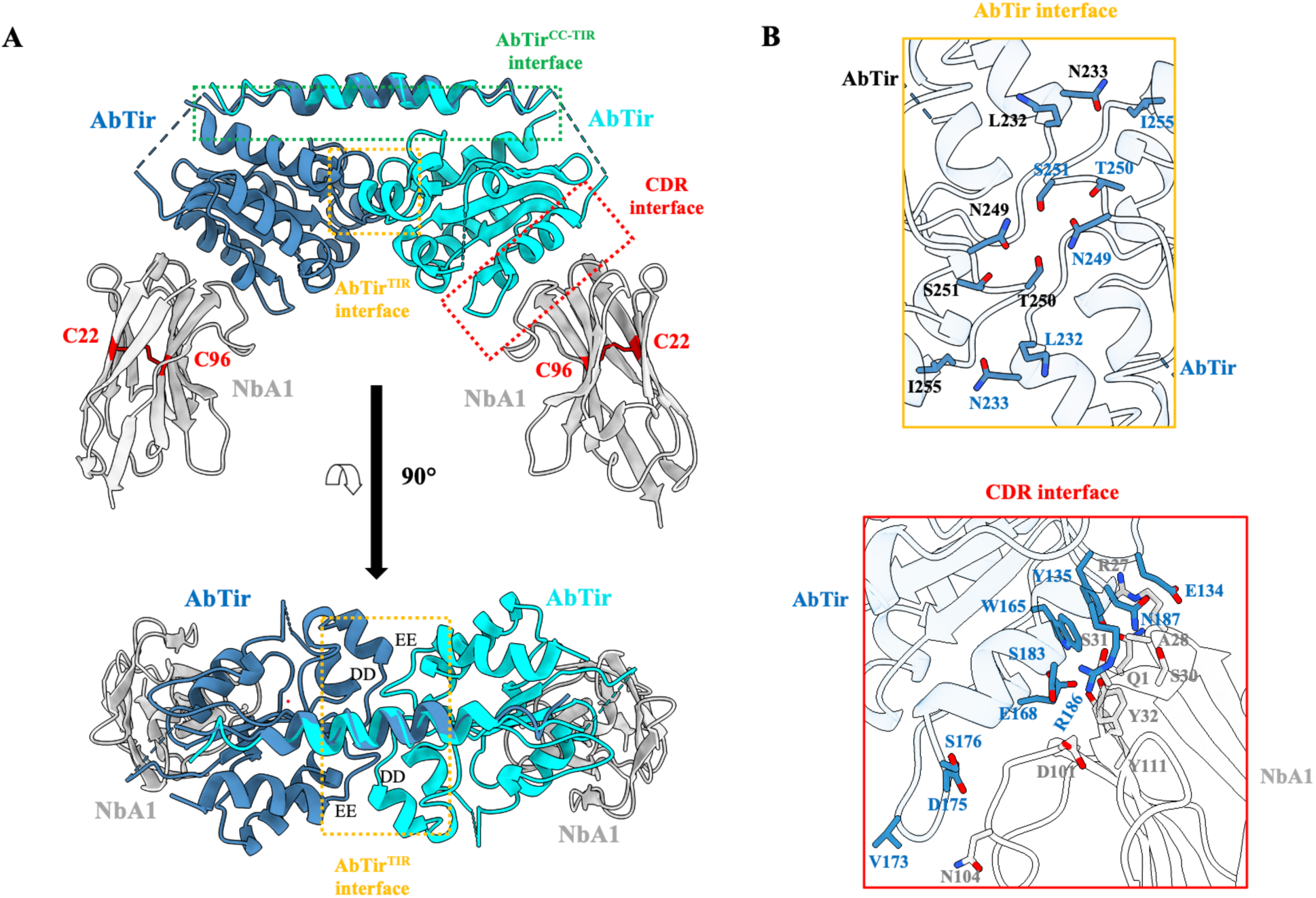
Crystal structure of AbTir in complex with NbA1. (**A**) Structure of the NbA1:AbTir complex. The most likely quaternary structure configuration in solution involves symmetry-related molecules within the crystals. The two AbTir protomers are shown as steel-blue and cyan cartoons. NbA1 is displayed as a grey cartoon. A disulfide bond between residues C22 and C96 is shown in red stick representation. (**B**) Interfaces observed in the AbTir:NbA1 complex. The AbTir interface is mediated by 6 hydrogen bonds. The CDR interface is mediated by 16 hydrogen bonds. Detailed hydrogen-bonding interactions are listed in Table 2.

**Table 1:** Crystallographic data collection and refinement statistics.

|  | AbTir: NbA1 | AbTir<sup>TIR-W204A</sup> | AbTir<sup>TIR-E208A</sup> |
| --- | --- | --- | --- |
| PDB ID | 26OL | 26MN | 26MQ |
| Space group | P4<sub>1</sub> 2 2 | P1 2<sub>1</sub> 1 | P 1 2<sub>1</sub> 1 |
| *a*, *b*, *c* (Å) | 71.45, 71.45, 164.46 | 55.52, 68.16, 72.78 | 54.55, 68.45, 71.80 |
| *α*, *β*, *γ* (°) | 90, 90, 90 | 90, 112.27, 90 | 90, 112.01, 90 |
| Resolution (Å) | 48.30 - 2.65 (2.78 - 2.65) | 47.91 - 2.55 (2.66 - 2.55) | 47.72 - 1.99 (2.04 - 1.99) |
| Total reflections | 312,953 (32,979) | 112,242 (11,973) | 124,204 (8,805) |
| Unique reflections | 23,516 (2,701) | 16,445 (1,811) | 32,647 (2,307) |
| Completeness (%) | 93.09 (79.22) | 98.7 (90.01) | 97.20 (95.00) |
| Multiplicity | 13.3 (12.2) | 6.8 (6.6) | 3.8 (3.8) |
| Wilson B-factor (Å<sup>2</sup>) | 45.90 | 46.32 | 21.72 |
| R<sub>meas</sub> | 0.26 (1.90) | 0.22 (1.78) | 0.07 (0.44) |
| R<sub>merge</sub> | 0.25 (1.90) | 0.20 (1.64) | 0.06 (0.38) |
| R<sub>pim</sub> | 0.07 (0.54) | 0.11 (0.93) | 0.04 (0.23) |
| Mean I/sigma(I) | 9.7 (1.3) | 4.8 (0.8) | 12.2 (2.6) |
| CC<sub>1/2</sub> | 0.996 (0.638) | 0.995 (0.612) | 0.998 (0.908) |
| **Refinement** |  |  |  |
| R<sub>work</sub> | 0.23 (0.29) | 0.21 (0.33) | 0.17 (0.24) |
| R<sub>free</sub> | 0.27 (0.37) | 0.27 (0.39) | 0.21 (0.30) |
| RMS bonds (Å) | 0.002 | 0.005 | 0.010 |
| RMS angles (°) | 0.49 | 0.908 | 1.11 |
| Ramachandran favoured (%) | 95.20 | 96.05 | 95.65 |
| Ramachandran outliers (%) | 0.70 | 0.38 | 0.19 |
| Rotamer outliers (%) | 0 | 0 | 0 |
| Clashscore | 2.3 | 7.4 | 3.6 |
| Average B-factor (Å<sup>2</sup>) | 52.83 | 55.20 | 28.63 |

1. The values in parentheses are for the highest-resolution shell. The statistics were calculated using Aimless and MolProbity.
2. $R_{\text{merge}} = \sum_{\text{hkl}} \sum_j |I_{\text{hkl},j} - \langle I_{\text{hkl}} \rangle| / (\sum_{\text{hkl}} \sum_j I_{\text{hkl},j})$ ; $R_{\text{meas}} = \sum_{\text{hkl}} [N/(N-1)]^{1/2} \sum_j |I_{\text{hkl},j} - \langle I_{\text{hkl}} \rangle| / (\sum_{\text{hkl}} \sum_j I_{\text{hkl},j})$ ; $R_{\text{pim}} = \sum_{\text{hkl}} [1/(N-1)]^{1/2} \sum_j |I_{\text{hkl},j} - \langle I_{\text{hkl}} \rangle| / (\sum_{\text{hkl}} \sum_j I_{\text{hkl},j})$ ; $R_{\text{work}} = \sum_{\text{hkl}} |F_{\text{obs}} - F_{\text{calc}}| / \sum |F_{\text{obs}}|$ ; $R_{\text{free}}$ is equivalent to $R_{\text{work}}$ , with 5% of data excluded from refinement process. $|F_{\text{obs}}|$ and $|F_{\text{calc}}|$ represent the observed and calculated structure factor amplitudes.

**Table 2:**
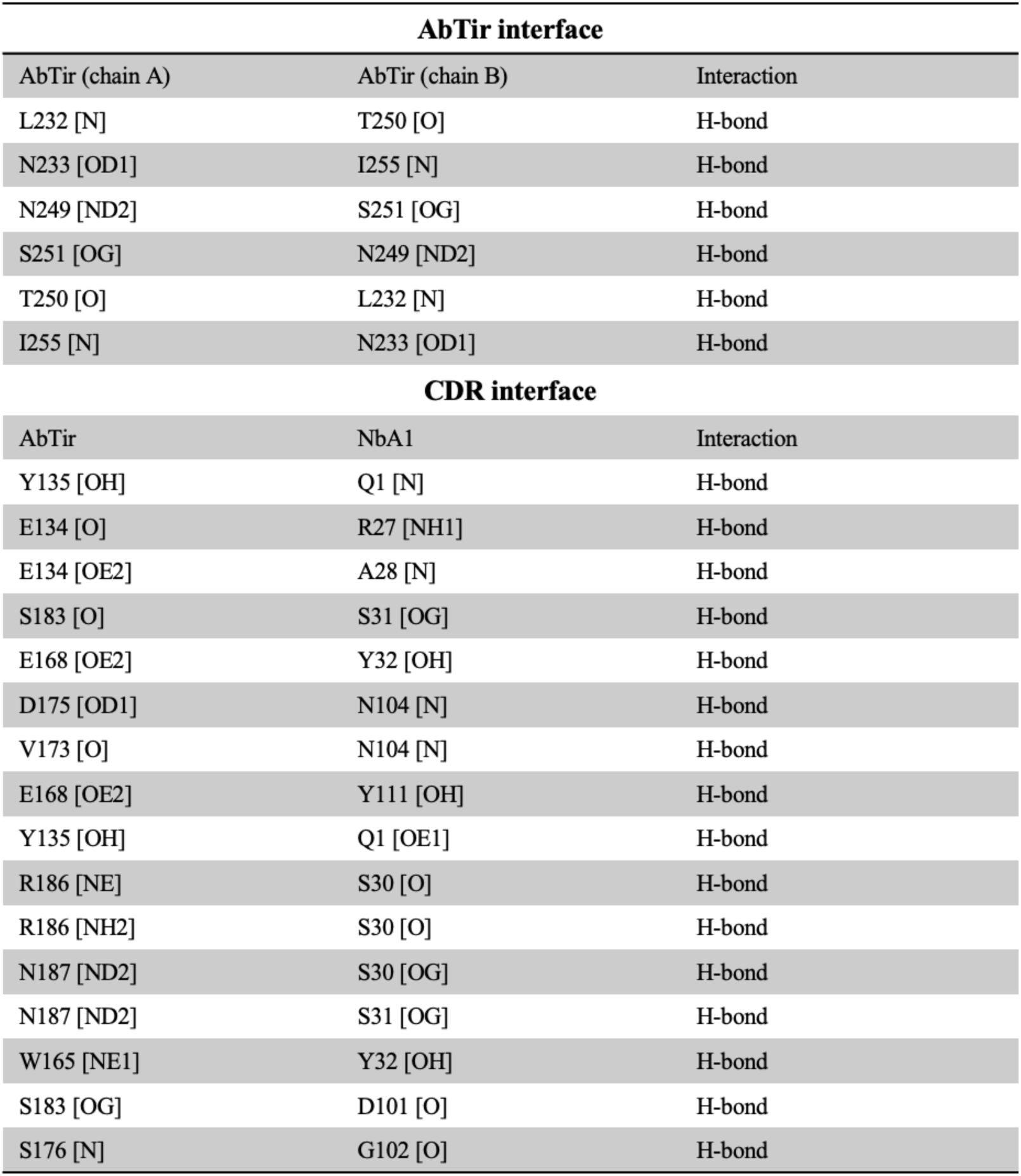
Interactions in the interfaces observed in the AbTir:NbA1 complex.

In the AbTir dimer, the primary dimerization interface (termed the “AbTir^TIR^ interface”) is formed between the two TIR domains and involves the DD and EE loops from each protomer, with a buried surface area (BSA) of 1,248 Å^2^ (Figure 2). This interface is mainly stabilized by six hydrogen bonds between residue pairs L232:T250, N233:I255, N249:S251 from opposing protomers (Figure 2B, Table 2). A secondary interface (termed the “AbTir^CC-TIR^ interface”) is formed between the CC-domain helix and the TIR-domain dimer (Figure 2A). Each CC helix buries approximately 178 Å² of surface area (BSA) against the opposing TIR domain and the interaction is mediated primarily through contacts with the βD and βE strands, the EE loop, and the αE helix in the TIR domain. This relatively small interface suggests that this interaction is likely to be weak and transient. The electron density for the CC-domain helix is poorly defined, preventing unambiguous model building. Consequently, residues 92 to 119 were modelled in two opposite orientations based on their approximate spatial relationship to the associated TIR domains. Future mutagenesis studies will be required to further validate and characterise this intermolecular CC:TIR interactions in AbTir.

The largest interface between AbTir and NbA1 corresponds to the complementarity-determining region (CDR) interface, with a BSA of approximately 1,251 Å^2^ (Figure 2, S2A, Table 2). This interface involves the CDR1 and CDR3 loops of NbA1 engaging the βB strand, BB-loop and αB helix of AbTir. Notably, this interface does not contribute to crystal packing within the asymmetric unit. Instead, crystal contacts are formed via the NbA1 framework regions FR2, FR3 and FR4 (the FR interface, Figure S2A), with a smaller BSA of 839 Å^2^. The CDR interface defines the binding specificity of NbA1 for AbTir and is mainly mediated by 16 hydrogen bonds between NbA1 and AbTir (Figure 2B). A disulfide bond between residues C22 and C96 is present in NbA1, contributing to stabilisation of the nanobody structure (Figure 2A).

### AbTir dimer in the complex represents the inactive state

The cryogenic electron microscopy (cryoEM) structure of AbTir^TIR^ bound to the NAD^+^ mimic 3AD (8-amino-isoquinoline adenine dinucleotide) showed a parallel, two-stranded filamentous assembly of TIR domains (PDB (Protein Data Bank): 7UXU)^8^. In this structure, 3AD is positioned between two AbTir^TIR^ molecules in a head-to-tail configuration, mediated by the BB-loop-containing “BE interface”, likely representing the active state of the protein^8^. By contrast, the AbTir:NbA1 crystal structure, determined in the present work, adopts a markedly different conformation, including a rotation of the BB-loop and αB helix by 50-55° (Figure 3A, B). The TIR domain in the AbTir:NbA1 complex closely resembles the previously reported crystal structure of an isolated AbTir TIR domain (PDB: 7UWG, 8G83)^8,28^, with a RMSD (root mean square deviation) value of 0.874 Å over 131 Cα atoms. A minor difference is observed in the BB-loop region, where NbA1 binding induces a ∼5° shift of the BB loop towards the nanobody, leading to the disorder of the WxxxE motif (residues 204 to 208, Figure 3B).

**Figure 3:**
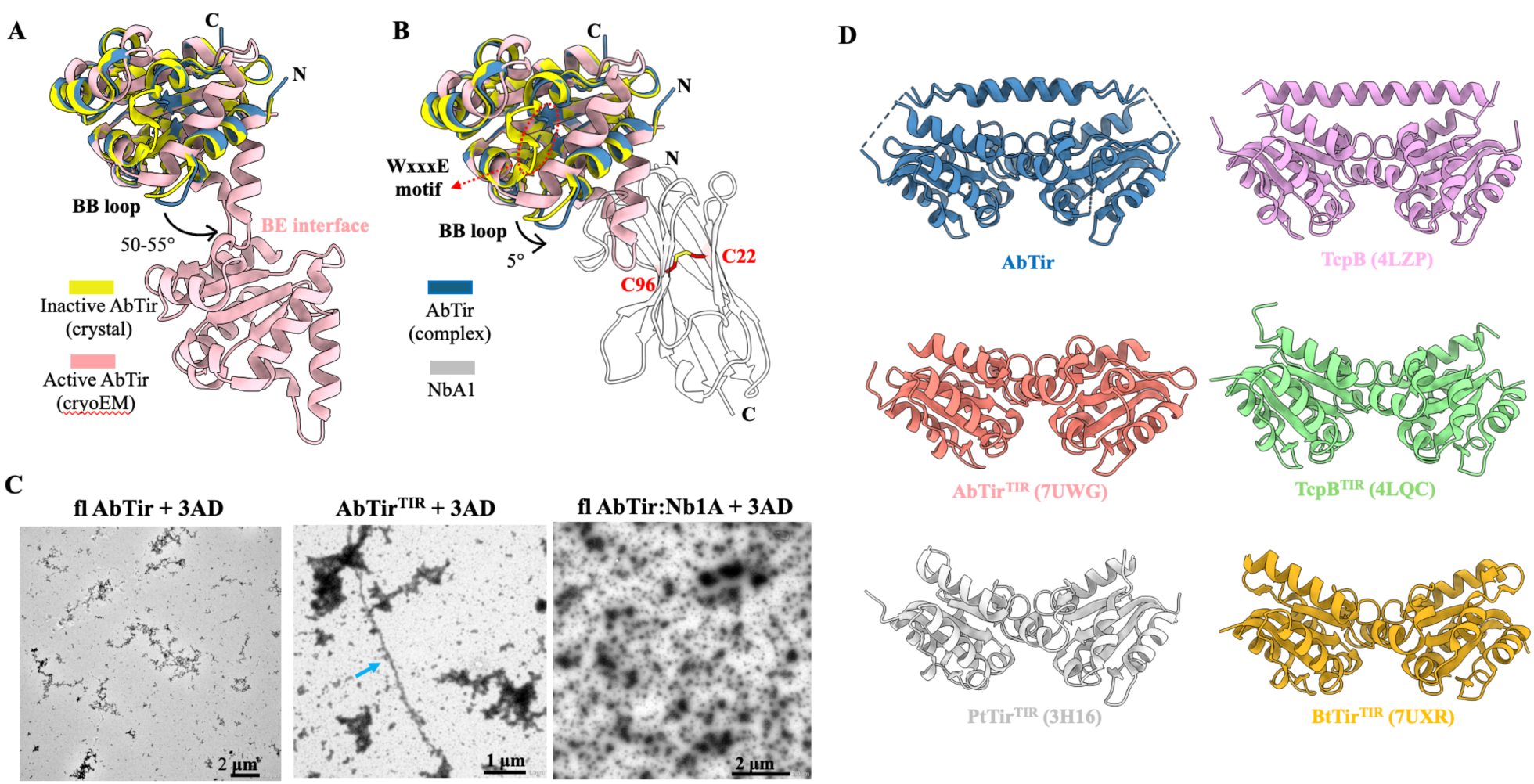
AbTir in complex with NbA1 represents the inactive conformation. (**A**) Structural superposition of AbTir^TIR^ (yellow, PDB:7UWG) and AbTir (steel-blue) from the NbA1 complex onto a protomer from the single head-to-tail chain of the 3AD-bound AbTir^TIR^ filament (pink, two protomers shown, PDB: 7UXU). In the active filament, the BB-loop and αB helix undergo a 50-55° rotation, promoting formation of the BE interface where the active site is located. (**B**) Structural superposition of AbTir^TIR^ (yellow, PDB:7UWG) and the protomer from the 3AD-bound AbTir^TIR^ filament (pink, PDB: 7UXU) onto AbTir (steel-blue) in the NbA1 complex. Binding of NbA1 induces a ∼5° rotation of the BB loop towards the nanobody, and results in disorder of the WxxxE motif in AbTir. (**C**) Negative-stain EM analysis of the AbTir:NbA1 complex incubated with 3AD. Unlike AbTir^TIR^ only, neither the complex nor full-length AbTir forms filaments in the presence of 3AD. (**D**) Available bacterial Tcp structures showing dimeric arrangements.

### Crystal structures of AbTir^TIR^ WxxxE-motif mutants resemble the inactive wild-type AbTir^TIR^

The WxxxE motif is conserved among most bacterial Tcps^32^ and is essential for NADase catalytic activity^8^. In particular, the tryptophan residue at position 204 (W204) is required for cyclization of ADPR into 2’cADPR, and NAD-RNA decapping activity^8,30^. The equivalent tryptophan is also conserved and important for enzyme activity in eukaryotic TIR proteins, including human SARM1 (sterile alpha and Toll/interleukin-1 receptor motif-containing 1; W638) and the flax L6 protein (W131)^33,34^. To further investigate the role of this motif, we determined the crystal structures of the TIR domains of the tryptophan mutant AbTir^TIR-W204A^, and the catalytic glutamate mutant AbTir^TIR-E208A^ (Figure S2C). Both structures closely resemble the monomeric wild-type AbTir^TIR^ crystal structure (PDB: 7UWG), with the RMSD values of ∼0.5 Å over 135 Cα atoms. All structures adopt the same symmetric dimer configuration, and none have the active sites formed (Figure S2C, 3D). Consistently, the catalytic residue E208 is also not in a position suitable for catalysis. These observations further confirm that the TIR-domain conformation in the AbTir:NbA1 complex likely represents the inactive state of the protein. In agreement, incubation of the AbTir:NbA1 complex with NAD^+^ analogue 3AD no longer resulted in the filamentous assemblies observed for the active AbTir^TIR^ in presence of 3AD (Figure 3C). As NbA1 binds directly to the BB-loop region, which undergoes substantial rearrangement between inactive and active conformations, these results further suggest that NbA1 preferentially recognizes and stabilizes the inactive state of AbTir.

### Single CC domain stabilizes the AbTir TIR-domain dimer

The CC domain of TcpB has previously been reported to be flexible and to possess only limited α-helical character^21,22^. Our crystal structure suggests that the CC domain of AbTir shares similar properties. Notably, only one single helix from the CC domain is observed and packs against the AbTir dimer. Modelling the CC helices from both protomers in opposite orientations indicates that either helix could geometrically engage this surface, while simultaneous binding of both helices would result in steric clashes. The helices therefore bind in a mutually exclusive manner, with one engaging the TIR-domain dimer, whereas the other is presumably disordered. These observations suggest that the CC domain acts as a dynamic dimerization module rather than forming the stable parallel or anti-parallel CC interaction more commonly associated with CC-mediated interactions^25^. This asymmetric arrangement further implies that a single CC helix is sufficient to stabilise the TIR:TIR dimer, consistent with biochemical studies showing that the CC domain is required for efficient oligomerization and regulation of NADase activity. Analogous to AbTir, a closely related arrangement is also observed in the crystal structure of *Brucella* TcpB (PDB: 4LZP, Figure 3D). In that study, the absence of density corresponding to one N-terminal CC helix was interpreted as possible evidence of proteolytic degradation during or prior to crystallization^21^. However, in the absence of direct evidence for degradation, we propose that TcpB may be in a similar mutually exclusive binding situation as observed in our structure. The observation of the analogous CC-binding mode in both TcpB and AbTir suggests that rather than being a crystallization artefact, the dynamic and intrinsically flexible nature of the CC domain may represent a conserved structural feature of this family of bacterial CC-TIR proteins.

### AbTir belongs to a defence-associated family of bacterial TIR domain-containing proteins

Many bacterial TIR domains participate in antiphage defence systems, where phage infection triggers abortive infection that limits phage spread through the death of infected bacterial cells^15,16,35,36^ To investigate whether AbTir also plays a similar role, we reconstructed a focused phylogeny of close AbTir homologs, using AbTir as the seed sequence (Figure 4A). Rooting the tree with ThsB from Thoeris type I systems revealed multiple well-supported clades that correlate with distinct domain architectures, including CC-TIR proteins, retron-associated TIRs and DUF1883-TIR proteins. To assess potential functional roles, we analyzed the genomic neighborhoods of these proteins and annotated whether they are associated with known defence systems. This information was then overlaid onto the phylogeny, revealing that defence-associated genomic contexts are prevalent across the entire tree. Consistently, defence scores were high across clades, including those previously proposed to encode translocated or virulence-associated effectors. This widespread enrichment and the presence of clades that constitute known defense systems (e.g. Retron XII) indicates that close homologs of AbTir are generally linked to defence-associated loci.

**Figure 4:**
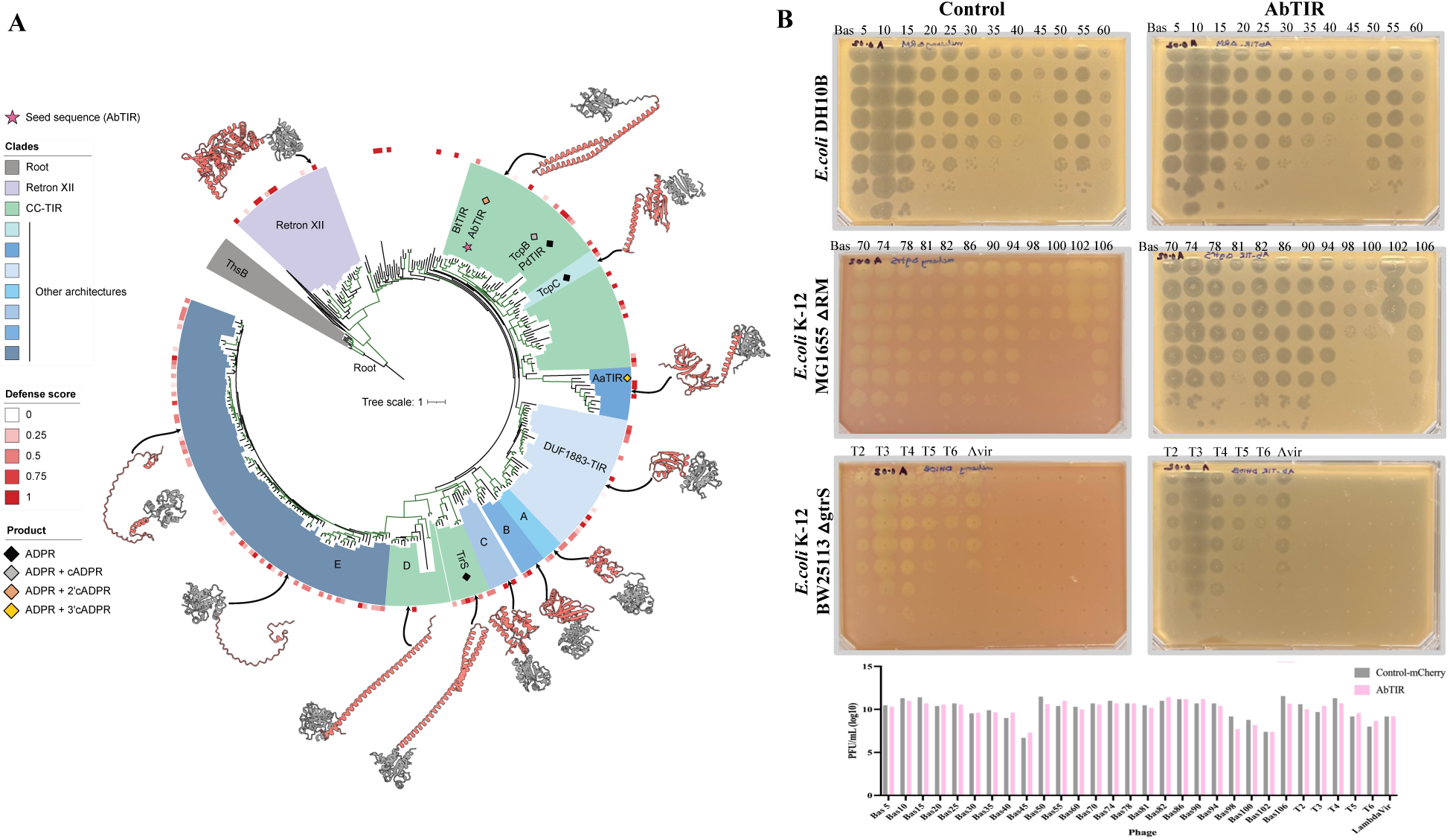
Analysis of possible antiphage defence functions of AbT*ir*. (**A**) Maximum-likelihood phylogeny of TIR domains homologous to AbT*ir*. The AbT*ir* seed sequence is indicated by a star. Major clades are highlighted and labeled based on tree topology and recurrent full-length domain architectures, including retrons, CC-TIRs, DUF1883-TIRs, and other architectures. The outer ring shows the defense score (red scale)^15^, reflecting enrichment of homologs in genomic neighborhoods that contain known anti-phage defense systems. Symbols indicate enzymatic products reported for previously characterized TIR proteins mapped onto the tree, including ADPR and cADPR variants. Representative AlphaFold 3 predictions are shown for selected clades, with the TIR domain colored in grey and additional domains in red. (**B**) Phage-spot assays testing the potential antiphage activity of AbTir in heterologous E. coli systems. E. coli DH10B, K-12 MG1655 ΔRM, and K-12 BW25113 ΔgtrS strains expressing AbTir were challenged with dsDNA phages from the *BASEL* phage collection, including T2, T3, T4, T5, T6, virulent Lambda and Bas phages. Serial dilutions of individual phage stocks were spotted onto bacterial lawns expressing either control vectors encoding mCherry or AbTir. The bar graph shows the phage titre determined from the spot assays. Data are from a single experiment (n = 1). Under the conditions tested, expression of AbTir did not result in detectable reductions in phage plaquing efficiency compared with the corresponding control strains, indicating no observable antiphage activity in the surrogate E. coli hosts.

Previously characterized bacterial TIR-domain enzymes distributed across the phylogenetic tree have been shown to cleave NAD⁺, generating nicotinamide and ADPR, with a subset additionally producing cyclic ADPR variants. The distribution of these activities across multiple clades is consistent with a conserved NAD⁺-processing function among bacterial TIR domains, while differences in reaction products appear to reflect diversification within a shared enzymatic framework. Structure predictions for representative proteins further reveal a conserved TIR core combined with diverse accessory domains, suggesting that functional diversification is primarily driven by regulatory and architectural variation, rather than changes in the catalytic mechanism. Together, these analyses place AbTir within a large and evolutionarily diverse family of bacterial TIR domain-containing proteins that are broadly associated with defence-related genomic contexts and share a conserved NAD^+^-processing core.

### AbTir shows no antiphage defence activity in the surrogate host *E. coli*

Functional characterization of defence systems in *A. baumannii* remains technically challenging, because genetic manipulation and expression systems in this organism are substantially less established than those available for model bacteria such as *E. coli*^37,38^. In addition, access to and distribution of *A. baumannii* isolates are often subject to biosafety regulation and restrictions associated with multidrug-resistant strains^39^. We therefore used a surrogate *E. coli* system to investigate the potential antiphage activity of AbTir. To assess the antiphage activity of AbTir, we challenged *E. coli* strains expressing AbTir from an arabinose-induced vector with a panel of 30 dsDNA phages, including T-phages and representatives spanning the major phage families present in the BASEL phage collection^40^. Under the conditions tested, AbTir expression did not result in measurable reductions in phage plaquing efficiency (Figure 4B). These results suggest that AbTir alone is insufficient in confering detectable antiphage activity in the surrogate *E. coli* system. Host-specific factors or regulatory mechanisms present in *A. baumannii* may be required for its physiological function.

## Discussion

This study provides insights into the molecular basis by which the CC domain regulates AbTir NADase activity. Previous studies on AbTir, using recombinantly produced protein, showed that the full-length protein exhibits an initial lag phase, before NAD⁺ nucleosidase activity increases rapidly^8^. By contrast, the isolated TIR domain lacks this lag phase and shows a higher activity. Similarly, full-length TcpB protein expressed in yeast displays lower NAD⁺ nucleosidase activity than its isolated TIR domain, indicating that the N-terminal CC region restrains enzymatic activity prior to activation^8,41^. By determining the crystal structure of the full-length AbTir in complex with the nanobody NbA1, we reveal a symmetric dimeric assembly in which a single CC-domain helix from one protomer interacts with both TIR domains in the dimers, thereby stabilizing the dimeric configuration. Together with the observed loss of activity of the AbTir:NbA1 complex and the inhibition of filament formation in the presence of 3AD, these findings demonstrate that the CC domain plays a central role in maintaining AbTir in an NADase-inactive dimeric state.

Comparison of the AbTir:NbA1 complex with the cryoEM structure of AbTir filaments (PDB: 7UXU), which represents the active state, reveals significant conformational differences in the BB-loop and αB helix regions. Upon converting to the active state, these two regions undergo substantial conformational changes while forming the active site at the BE interface (Figure 3A). By contrast, the AbTir TIR domain in our nanobody-bound complex closely resembles the crystal structures of AbTir^TIR^ (PDB: 7UWG, 8G83)^8,28^. It also resembles the crystal structures of other bacterial TIR proteins including TcpB (4LZP, 4LQC, 4C7M)^7,21,22^, PdTir (PDB: 3H16)^42^ and BtTir (PDB: 7UXR)^8^. In these structures, the BB-loop and αB helix adopt a conformation incompatible with active-site formation (Figure 3D). In addition, a symmetric dimeric assembly similar to that observed in our complex is conserved across these bacterial TIR crystal structures (Figure 3D). In all these structures, the active sites required for NAD^+^ cleavage are not formed, and the catalytic glutamate is positioned in a location incompatible with catalysis. Notably, although many human TIR domains associated with TLR and interleukin-1 receptor signalling contain a conserved glutamate residue at an analogous sequence position, available structures indicate that the conserved glutamate is also positioned in a location incompatible with catalysis. Recombinantly produced TIR domains from these human proteins generally lack detectable NADase activity, despite reports of NADase activity in some cell lysate-based assays^5,9,33,43^.

The conserved WxxxE motif is essential for NADase activity, with the tryptophan residue required for both NAD^+^ cleavage and generation of cADPR variants, while the glutamate residue functions directly in catalysis. In some plant TIR domains, the equivalent motif has diversified to support alternative activities, including DNA and RNA binding and cleavage, as well as cyclic nucleotide synthesis^34^. In these proteins, a cysteine residue important for cAMP (cyclic adenosine monophosphate) and cGMP (cyclic guanosine monophosphate) synthetase activity (e.g. C132 in the flax L7 protein), replaces the threonine found in the AbTir WxxxE motif (T205)^34^. Consistently, AbTir is unable to bind DNA and RNA and thus it is unlikely to cleave them and synthesise cyclic nucleotide monophosphates^34^. These observations underscore the functional versatility of the WxxxE motif and its central role in determining the biochemical activity of TIR domains. In concurrence, the WxxxE motif is disordered in the crystal structures of AbTir:NbA1 and many of bacterial Tcps, in line with these structures representing inactive forms. Together, these observations support the notion that bacterial TIR crystal structures generally capture an inactive dimeric conformation.

CC domains are versatile structural modules that participate in a wide range of biological processes including protein oligomerisation, signal transduction and membrane targeting^24^. In most characterised proteins, CC domains typically mediate protein assembly through the formation of parallel or anti-parallel helical bundles^25,44^. In some proteins, CC domains also function as intramolecular regulatory elements, by interacting with other domains, thereby stabilising autoinhibited conformations prior to activation^45,46^. Our data suggest that AbTir employs a distinct and unusual strategy in which only a single CC helix contributes to stabilization of the inactive TIR dimer. Structural analysis indicates that only one CC helix can be accommodated at the observed interface, such that occupation by one CC sterically excludes the symmetry-related CC from adopting an equivalent position. Consequently, the second helix must find a different position in the structure, and considering we cannot visualize it in the crystal, we assume it is disordered. This mechanism represents a form of symmetry breakage driven by the mutually exclusive occupation of overlapping structural positions. We developed a Cα-based pairwise distance scanning pipeline to search the PDB and identified a limited number of protein structures showing a similar alternate-conformation behaviour to the one observed in the AbTir structure. In procaspase-1, linker segments from two symmetry-related protomers compete for occupancy of a common cavity at the dimer interface, suggesting that simultaneous occupation of both positions is sterically prohibited (PDB: 3E4C)^47^. Similarly, galactose-specific legume lectins from *Dolichos biflorus* (PDB: 1LU1)^48^ and *Dolichos lablab* (PDBs: 3UJO, 3UJQ, 3UK9 and 3UL2)^49^ exhibit static disorder of a central α-helix within an otherwise symmetric tetrameric assembly, indicating that equivalent structural elements cannot be accommodated simultaneously within the oligomeric core^48,49^. In none of these cases, the functional relevance of the phenomenon has been explained.

In AbTir, this asymmetric arrangement likely reflects a functional compromise between stability and flexibility. A single CC helix provides sufficient contact to hold the two AbTir protomers together, thereby promoting dimer formation, but at the same time, remains dynamic enough to disengage when conformational rearrangements are required for activation. Given that AbTir undergoes substantial structural changes when forming the active filamentous structure, particularly around the catalytic interface^8,28^, excessive stabilization by a canonical parallel or anti-parallel CC bundle would likely constrain these transitions and retain the protein in an inhibited state. Thus, the single-helix CC interaction may act as a tunable regulatory element that stabilizes the inactive dimer, while allowing controlled release of inhibition upon activation cues. As a similar arrangement has also been observed in TcpB, we propose that the CC domain of AbTir and presumably CC domains in other bacterial CC-TIR proteins may inhibit the nucleosidase activity by stabilizing an inactive dimeric configuration and preventing the head-to-tail TIR association required for the formation of the active site, while still allowing inhibition to be released when conformational changes are triggered during activation.

The mutually exclusive location of only one helix could perhaps allow the second helix to play a separate functional role. Bacterial CC-TIR proteins can also function as virulence factors by mimicking host TIRs^8,9^. TcpB from *Brucella melitensis*, for example, has been shown to mimic MAL (a TLR adaptor protein) by binding to the plasma membrane using its N-terminal CC domain, thereby suppressing MAL-dependent TLR signalling^22,26^. This observation suggests that CC-domain helices may serve additional roles beyond NADase activity regulation, including membrane anchoring and subcellular localization. While direct evidence of AbTir-modulated host innate immune signalling is lacking, the asymmetric arrangement of its CC domain raises the possibility that the two CC helices may adopt functionally distinct roles, for example membrane association or host-factor engagement.

Bacterial TIR proteins constitute a widespread and mechanistically diverse class of immunity-associated proteins involved in antiphage defence^36^. Our phylogenetic analyses place AbTir within a family of bacterial TIR-domain proteins that are broadly linked to defence-related genomic contexts. To investigate if AbTir exhibits antiphage activity, we performed phage-spot assays. Given the technical challenges associated with genetic manipulation and phage infection in *A. baumannii*^37,38^, we employed the model organism *E. coli* as a tractable heterologous system for initial functional testing. However, phage-spot assays using the *E. coli* system did not reveal detectable antiphage activity under the conditions tested. This apparent discrepancy may reflect the highly context-dependent nature of bacterial defence systems. Antiphage pathways exhibit strong host and phage specificity and often require particular signalling partners, accessory proteins, regulatory mechanisms, or phage-derived activation signals to initiate defence responses^36,50^. *A. baumannii* and *E. coli* differ substantially in their cellular physiology, envelope composition, signalling pathways, and phage-susceptibility profiles, all of which may influence defence-system activities. Indeed, heterologous expression systems may fail to fully recapitulate the native regulatory and activation context required for *A. baumannii* immune pathways. Therefore, the absence of detectable protection in *E. coli* does not exclude the possibility that AbTir functions in bacterial defence within its native or a more closely related host background. Future studies using *A. baumannii* or related species, together with cognate phages and potential accessory factors, will be required to fully evaluate the physiological role of AbTir in bacterial immunity.

To conclude, our structural and biochemical analyses reveal the unusual mechanism used by the CC domain to stabilize an inactive AbTir dimer and prevent formation of the catalytic interface. This architecture provides a simple mechanism by which bacterial TIR proteins may maintain their NADase activity in a suppressed state, until activation is triggered. The observation that related dimeric assemblies are present across multiple bacterial TIR domains suggests that such negative regulation may be broadly conserved among bacterial TIR proteins. At the same time, the occurrence of structurally distinct symmetric dimers in several mammalian TIR structures^17,33,51^ hints that different TIR families may have evolved distinct structural solutions to achieve analogous regulatory outcomes. Future studies should aim to determine how the CC domains contribute to AbTir function in cells, including whether the two CC helices adopt distinct regulatory and localisation-related roles. In addition, further investigation using native or closely related bacterial hosts, together with cognate phages and potential accessory components, will be required to clarify the physiological role of AbTir in bacterial antiphage defence and cellular signalling.

### Experimental Procedures

#### Cloning

The cDNA of AbTir (GenBank: EXB04249.1; residues 1-267), purchased from Integrated DNA Technologies, was amplified using AccuPower® Pfu PCR PreMix (Bioneer Pacific). The amplified product was cloned into the pMCSG7 plasmid at the SSpI site, using the ligation-independent cloning method^52^. Initial candidates of nanobodies were selected from an alpaca immunized with the purified AbTir protein. Immunisation and handling of the alpaca for scientific purposes was approved by Agriculture Victoria, Wildlife & Small Institutions Animal Ethics Committee, project approval No. 26-17. Following immunization, blood was collected for the preparation of lymphocytes. The total alpaca lymphocyte mRNA was extracted and amplified by RT-PCR, then cloned into a phage-display vector to generate a nanobody library. After phage display selection against immobilized AbTir, individual clones were screened by enzyme-linked immunosorbent assay (ELISA) for binding to recombinant His-tagged AbTir. Bound nanobodies were detected using an anti-His-tag antibody. Positive clones were sequenced, and candidates with unique CDR3 regions were selected for further analysis. The plasmids encoding AbTir^TIR^ mutants were generated using the Q5® Site-Directed Mutagenesis Kit purchased from New England BioLabs (NEB). The pure plasmids were prepared using QIAprep Spin Miniprep Kit (Qiagen) and the sequences were confirmed using Sanger sequencing by the Australian Genome Research Facility (Brisbane).

#### Protein expression

The plasmids of AbTir or AbTir TIR-domain mutants were transformed into BL21-Gold (DE3) competent cells (Agilent Technologies, Inc.), which were grown on LB (lysogeny broth) agar plates containing 100 μg/mL ampicillin at 37 °C. The next day, a 10 mL starter culture was grown for 16 hours at 37 °C in LB medium containing 100 μg/mL ampicillin. The following day, 1 mL of the 16-hour culture was added to 1 L autoclaved LB-ampicillin (100 μg/mL) media in 2.5 L ultra-yield flasks (Thomson’s Ultra Yield FlasksTM, Genesearch). The flasks were incubated at 37 °C in a shaking incubator (New Brunswick^TM^ Innova® 44) at 225 rpm until OD_600nm_ reached 0.6-0.8. IPTG (isopropyl β-D-1-thiogalactopyranoside) (Merck, Millipore) was then added to a final concentration of 1 mM. The cells were incubated at 15 °C for 14-16 hours.

The nanobody constructs were transformed into WK6 competent cells on LB-ampicillin (100 μg/mL) plate. Nanobodies were expressed in TB (terrific broth) media, using the IPTG-induction method similar to the one used for the AbTir protein, and grown at 28 °C for 12-16 h.

#### Protein purification

To purify the AbTir protein and its mutants, the cells were harvested by centrifugation at 4000 × *g* (Beckman Coulter J-26 XPI, JLA 9.1 rotor) for 20 minutes at 4 °C. The supernatant was discarded, and the cell pellet was resuspended using ice-cold lysis/wash buffer (2 × PBS, 300 mM NaCl, 30 mM imidazole, 1 mM PMSF (phenylmethanesulfonylfluoride)). Bacterial cells were lysed using sonication (Branson, 10 s pulse, 10 s off at 40% amplitude). Lysed samples were then centrifuged at 15,000 × *g* (Beckman Coulter J-26 XPI, JA 20 rotor) for 40 minutes at 4 °C to remove the cell debris, and the supernatant was loaded onto a 5 mL HisTrap column (GE Healthcare) at 4 mL/min. After that, the column was washed using 20 column volumes (CVs) of ice-cold lysis/wash buffer. The protein was eluted using 10 CVs of elution buffer (100 mM HEPES pH 8.0, 500 mM NaCl, 500 mM imidazole). The eluted samples were analyzed using 15% SDS-PAGE, and the fractions containing pure proteins were pooled and dialyzed for 30 minutes in dialysis buffer (2 × PBS, 1 mM DTT) at 4 °C to remove imidazole. After 30 minutes, TEV (tobacco etch virus) protease was added and incubated overnight at 4 °C to remove the His-tag. The next day, dialyzed samples were passed through a 5 mL HisTrap column (GE Healthcare) to remove the TEV protease. Then, the flow-through was further purified with size exclusion chromatography using the S75 HiLoad 26/600 column (GE Healthcare) pre-equilibrated with the gel-filtration buffer (10 mM HEPES pH 8.0, 150 mM NaCl).

To purify the nanobodies, the cells were harvested by centrifugation at 5000 × *g* (Beckman Coulter J-26 XPI, JLA 9.1 rotor) for 10 minutes at 4 °C. The supernatant was discarded, and the cell pellet was resuspended using ice-cold resuspension buffer (1 × PBS, 20% w/v sucrose, 20 mM imidazole, 150 mM NaCl). The resuspended cells were incubated on ice for 15 minutes. 0.5 M EDTA (pH 8.0) was added to the resuspended cells to a final concentration of 5 mM and the cells were incubated for 20 minutes on ice. To prevent nickel’s chelation by EDTA, MgCl_2_ was added to a final concentration of 10 mM. Cell suspensions were then centrifuged at 20,000 × *g* (Beckman Coulter J-26 XPI, JA 20 rotor) for 30 minutes at 4 °C to remove the cell debris, and the supernatant was loaded into a 5 mL HisTrap column (GE Healthcare) pre-equilibrated with the buffer of 1 × PBS, 100 mM NaCl and 20 mM imidazole. After that, the column was washed using 5 CVs of 5% elution buffer (1 × PBS, 100 mM NaCl, 400 mM imidazole, pH 7.5) and ice-cold 95% wash buffer (1 × PBS, 100 mM NaCl, 20 mM imidazole). The protein was eluted using a linear gradient of 100% elution buffer. The eluted samples were analyzed using 15% SDS-PAGE, and the fractions containing pure proteins were pooled and concentrated. The sample was further purified with SEC using the S200 HiLoad 16/600 column (GE Healthcare) pre-equilibrated with the gel-filtration buffer (10 mM Hepes pH 7.5, 150 mM NaCl).

For the purification of protein-nanobody complexes, AbTir and nanobodies were mixed (1:2 ratio) and incubated at 4 °C overnight. The next day, the mixture was further purified with SEC using the Superdex 200 Increase 10/300 column (GE Healthcare) pre-equilibrated with the gel-filtration buffer.

#### NAD^+^ nucleosidase assays

1, N^6^-ethenoNAD (εNAD) (Sigma-Aldrich), a fluorescent analog of NAD^+^, was used as the substrate in the fluorescence-based NAD^+^ nucleosidase assays. 100 μM protein and 100 μM substrate were used. The assay was carried out in 96 well microplates (Greiner). Fluorescence intensity was measured using CLARIOstar® microplate reader (excitation wavelength 310-330 nm; emission wavelength 390-410 nm, readings every 1-3 minutes over 4 hours at 25 °C). The change in fluorescence over time was calculated from the slopes of the linear component of the curves.

#### SEC-MALS

A DAWN HELEOS II 10-angle light-scattering detector coupled with an Optilab rEX refractive index detector (Wyatt Technology), combined with Superdex 200 5/150 Increase size-exclusion column (GE Healthcare) connected to a Prominence HPLC (Shimadzu), was used for SEC-MALS analysis. The column was equilibrated with the gel filtration buffer for the complex, and 30 μL of 5 mg/ml protein-nanobody complex was run through the column at 0.25 mL/min. Molecular masses were calculated using Astra 6.1 (Wyatt Technology).

#### Negative-stain EM analysis

AbTiR^TIR^, full-length AbTir, and AbTir:NbA1 complex were diluted to 5 mg/mL and incubated with 2 mM 3AD at 25°C for 1 h. The protein samples were then diluted to 1 mg/mL in gel-filtration buffer containing 2 mM 3AD. 6 μL of the sample was loaded and incubated on the glow-discharged, carbon-coated copper grids for 5 minutes. The grids were then washed with the gel-filtration buffer containing 2 mM 3AD. The grids were stained with 2% uranyl acetate for 30 seconds twice and air-dried. The images were taken using JEOL JEM-1400 Flash electron microscope at 120 kV at various magnifications.

#### Protein crystallization

Proteins were crystallized using the hanging drop vapour diffusion method. Initial trays were set up using a Mosquito® Crystal instrument (SPT Labtech). Several initial hits were obtained for different protein samples within 1-3 days in different commercial crystallization screens. Diffraction-quality crystals of NbA1:AbTir complex were obtained in 0.1 M Bis-Tris propane pH 6.5, 0.2 M potassium sodium tartrate tetrahydrate, 25% PEG 3350 at 20 °C; and for AbTir^TIR-W204A^ and AbTir^TIR-E208A^, in 0.1 M Bis-Tris pH 5.5, 0.2 M lithium sulfate, 25% PEG 3350 at 20 °C.

#### Diffraction data collection and processing

The crystals were cryo-protected using the original well solution with additional 30% glycerol. The harvested crystals were immediately flash-cooled into liquid nitrogen. X-ray diffraction data were collected at the wavelength of 0.9537 Å at the Australian Synchrotron MX2 beamline ^53^ and indexed and integrated using XDS ^54^. Data scaling was performed using AIMLESS ^55^. Crystal structures were solved using molecular replacement (AbTir^TIR^ [PDB ID: 7UWG] and a nanobody [PDB ID: 5LHR] as search models) with Phaser ^56^. Model building and structure refinement were performed using Coot ^57^ and Phenix refine ^58^, respectively.

#### Identification of AbTir homologs

The AbTir protein sequence (UniProt A0A009IHW8) was used as a seed. Close homologs were retrieved from the RefSeq protein database using BLASTp (coverage >40%, sequence identity >50%), then dereplicated at 95% sequence identity (n = 10). These sequences were aligned with MAFFT (G-INS-i), and the TIR domain was extracted to build a profile HMM (pHMM). The pHMM was used to search RefSeq proteins for remote homologs using hmmsearch. Hits were retained with E-value < 1e-5 and additionally filtered by bitscore > 40 (threshold chosen based on the distribution of hit scores). This procedure yielded 1,844 proteins. Sequences were clustered with MMseqs2 at 80% sequence identity and 80% coverage, resulting in 502 protein families. For phylogenetic reconstruction, TIR domains were extracted based on the hmmsearch alignment coordinates.

#### Phylogenetic reconstruction and clade definition

Extracted TIR domains were aligned using MAFFT (G-INS-i). Alignment columns with >95% gaps were removed. A set of TIR domains from ThsB proteins (Thoeris Type I) was added as a distant homolog reference and used as an outgroup for rooting. The final alignment was used to infer a maximum-likelihood phylogeny with IQ-TREE3 using 1,000 ultrafast bootstrap replicates (-B 1000) and model search over WAG and Q.pfam (-mset WAG,Q.pfam). The tree was visualized in iTOL, and clades were manually annotated based on phylogenetic structure together with domain architectures of the corresponding full-length proteins.

#### Structural and functional annotation

Representative sequences from selected clades were submitted to AlphaFold 3 for structure prediction^59^ and mapped onto the tree. In structure renderings, the TIR domain is shown in grey and additional domains in red. Previously characterized TIR-domain proteins were mapped onto the phylogeny and annotated by reported reaction products. All mapped enzymes cleave NAD⁺ to generate nicotinamide and ADPR; some additionally produce cADPR. When a specific cADPR variant was experimentally determined, it is annotated accordingly.

#### Identification of alternate conformations in the PDB

To systematically identify structures in the PDB exhibiting alternate conformations analogous to the CC-domains in AbTir, we developed a Cα-based pairwise distance scanning pipeline. A local mirror of the PDB comprising 254,587 structures in mmCIF format, accessed on 1 April 2026, was scanned. Structures deposited as multi-model ensembles, characteristic of NMR experiments, were excluded from the analysis. For each remaining structure, the three-dimensional coordinates of all Cα atoms were extracted across all polymer chains using Biopython. A spatial index was constructed using the cKDTree implementation from scipy, and all Cα atom pairs within a distance threshold of 1.0 Å were identified. For pairs within the same chain, a sequence separation filter was applied such that only pairs with a residue index difference greater than 10 were retained, excluding trivially adjacent backbone atoms. Cross-chain pairs were included without sequence separation filtering. Flagged pairs were classified as intra-chain or inter-chain, based on whether the two residues belonged to the same or different polypeptide chains, respectively. The pipeline was validated against PDB entry 8P0D, a crystal structure of human 14-3-3σ in complex with a di-phosphorylated MDM2 peptide, which represents a known case of two overlapping peptide binding states modelled within a single chain at 0.5 occupancy each; this entry was correctly recovered by the scan. Using this approach, we identified a limited number of structures having alternate conformations similar to that observed in AbTir structure.

#### Defense score

Defense scores were computed using a procedure adapted from Doron et al.^15^. For each MMseqs2 family (80% identity and 80% coverage), we examined the genomic neighborhood (±10 genes) of each member and recorded whether at least one known defense system (detected with DefenseFinder) was present. The family-level defense score corresponds to the fraction of members with at least one defense system in their neighborhood (averaged across family members).

#### Phage isolation

A panel of 30 dsDNA phages, representative of the major phage families within the BASEL phage collection, was used in this study; the phages included T2, T3, T4, T5, T6, virulent Lambda, Bas5, Bas10, Bas15, Bas20, Bas25, Bas30, Bas35, Bas40, Bas45, Bas50, Bas55, Bas60, Bas70, Bas74, Bas78, Bas81, Bas82, Bas86, Bas90, Bas94, Bas98, Bas100, Bas102 and Bas106 ^40^. Phage stocks were amplified by inoculating 0.01% of phage lysate acquired from the Harms lab, into logarithmic phase *E. coli* cultures: *E. coli* DH10B, *E. coli* K-12 MG1655 ΔRM, and *E. coli* K-12 BW25113 ΔgtrS. The media were supplemented with 10 mM MgSO_4_ and 10 mM CaCl_2_. After overnight agitation at 37°C, cultures were centrifuged, and the supernatant was treated with 0.1 v/v chloroform. The supernatant was then passed through a 0.22 μm filter to obtain a clear phage lysate. Phages were tittered using the double-layer agar method.

#### Phage-spot assay

Competent strains of different *E. coli* hosts (*E. coli* DH10B, *E. coli* K-12 MG1655 ΔRM and *E. coli* K-12 BW25113 ΔgtrS) were transformed with the pBAD strains carrying the TIR domain-containing systems. Glycerol stocks were made and used to inoculate overnight cultures. The BASEL phage screen was performed on a medium-throughput scale using the Hamilton Microlab NIMBUS. Rectangular double-layer agar plates (Nunc™ OmniTray™ Single-Well Plate) were prepared, with 2 mL of overnight induced culture in the top agar (0.75%), supplemented with 100 μg/mL ampicillin and 0.02% L-arabinose. The base hard agar was supplemented likewise. In a 96-well plate, a dozen phages were serially diluted (10^-1^ to 10^-8^) across manually, of which 2 μL from each well was spotted on the double-layer agar plates using the liquid handler. The plates were imaged after overnight incubation at 37°C and the fold inhibition was noted manually.

## Acknowledgements

We acknowledge the use of the University of Queensland Remote Operation Crystallization and X-ray Diffraction (UQ-ROCX) facility at the Centre of Microscopy and Microanalysis (CMM) and the Australian Synchrotron MX beamlines at the Australian Nuclear Science and Technology Organisation (ANSTO), and thank the facility staff for their support. This work was supported by the National Health and Medical Research Council (NHMRC) (Investigator grants 2025931 to B.K. and 2016908 to W.H.T.); Australian Research Council (ARC) Laureate Fellowship (FL180100109 to B.K.); Snow Medical Research Foundation (SMRF2021-276 to G.J.K. and J.D.S.); ERC Starting Grant (PECAN 101040529), Vallee Scholar award, core funding from the Institut Pasteur and EvoCure consortium that receives financial support from Agence Nationale de la Recherche (ANR-24-RRII-0005) to M.H. and A.B.; and UQ RTP scholarship to M.K.M. The funders were not involved in the design of the study, data collection, analysis, interpretation of the data, the writing, or the decision to submit the article for publication.

---

**Figure S1:**
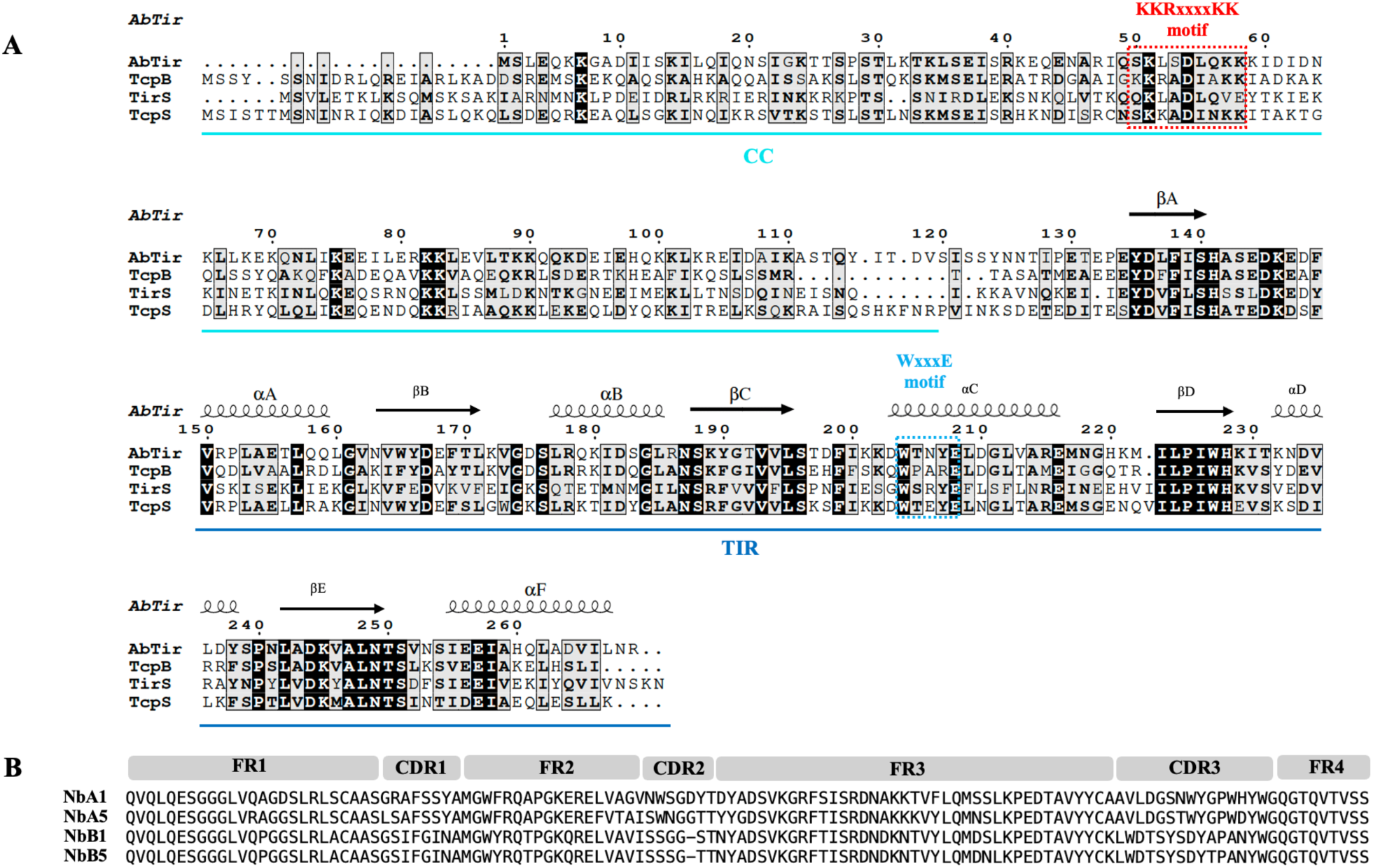
Sequences of bacterial TIR domain-containing proteins and nanobodies. (**A**) Sequence alignment of the representative Tcps including AbTir from *Acinetobacter baumannii*, TcpB from *Brucella melitensis*, TcpS from *Salmonella* Enteritidis and TirS from *Staphylococcus aureus*. Elements of secondary structure are indicated above the alignment, with ‘coils’ indicating α-helices and ‘arrows’ indicating β-strands. Conserved residues are highlighted in black. (**B**) Sequence alignment of nanobodies NbA1, NbA5, NbB1 and NbB5. The architecture of nanobodies is indicated above the alignment. FR: framework region; CDR: complementarity-determining region.

**Figure S2:**
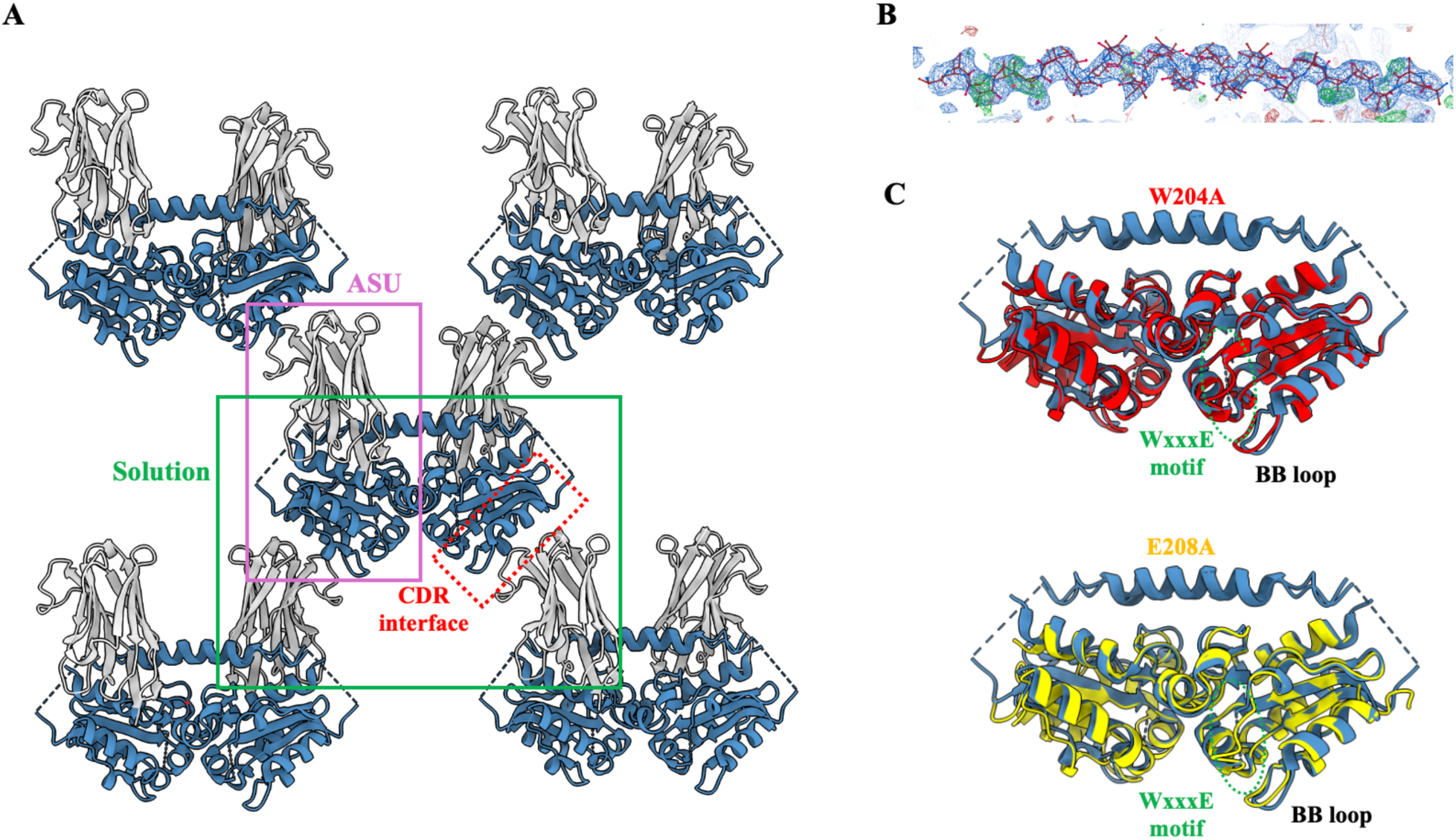
Crystal structures of AbTir and its mutants. (**A**) Packing of the crystals of AbTir in complex with NbA1. The AbTir:NbA1 complex in the ASU is highlighted with a purple box; the most likely AbTir:NbA1 complex in solution is highlighted with a green box. (**B**) Electron density map of the CC-domain helix. The 2Fo–Fc map is shown as a blue mesh and contoured at 0.7 RMSD. The refined model is shown in red stick representation. (**C**) Crystal structures of AbTir^TIR-W204A^ and AbTir^TIR-E208A^. AbTir dimer from the NbA1 complex is superimposed onto the crystal structures of the two mutants. Crystals of AbTir^TIR-W204A^ and AbTir^TIR-E208A^ mutants contain four molecules in the ASU. AbTir^TIR-W204A^ and AbTir^TIR-E208A^ are shown as red and yellow cartoon, respectively.

